# DNMT1 in Six2 progenitor cells is essential for transposable element silencing and kidney development

**DOI:** 10.1101/359448

**Authors:** Szu-Yuan Li, Jihwan Park, Kiwung Chung, Rojesh Shrestha, Matthew B Palmer, Katalin Susztak

## Abstract

Cytosine methylation (5mC) plays a key role in maintaining progenitor cell self-renewal and differentiation. Here, we analyzed the role of 5mC in kidney development by genome-wide methylation, expression profiling, and by systematic genetic targeting of DNA methyltransferases (Dnmt) and Ten-eleven translocation methylcytosine hydroxylases (Tet).

In mice, nephrons differentiate from Six2+ progenitor cells, therefore we created animals with genetic deletion of Dnmt1, 3a, 3b, Tet1, and Tet2 in the Six2+ population (Six2^Cre^/Dnmt1^flox/flox^, Six2^Cre^/Dnmt3a^flox/flox^, Six2^Cre^/Dnmt3b^flox/flox^, Six2^Cre^/Tet2^flox/flox^ and Tet1-/-). Animals with conditional deletion of Dnmt3a, 3b, Tet1 and Tet2 showed no significant structural or functional renal abnormalities. On the other hand, Six2^Cre^/Dnmt1^flox/flox^ mice died within 24hrs of birth. Dnmt1 knock-out animals had small kidneys and significantly reduced nephron number. Genome-wide methylation analysis indicated marked loss of methylation mostly on transposable elements. RNA sequencing detected endogenous retroviral (ERV) gene transcripts and early embryonic genes. Increase in levels of interferon (and RIG-I signaling) and apoptosis (Trp53) in response to ERV activity likely contributed to the phenotype development. Once epithelial differentiation was established, loss of Dnmt1, 3a, 3b, Tet1 or Tet2 in glomerular epithelial cells did not lead to functional or structural differences at baseline or following toxic glomerular injury.

Genome-wide cytosine methylation and gene expression profiling showed that Dnmt1-mediated DNA methylation is essential for kidney development by preventing regression of progenitor cells into a primitive undifferentiated state and demethylation of transposable elements.

**Significance:** Cytosine methylation of regulatory regions (promoters and enhancers) has been proposed to play a key role in establishing gene expression and thereby cellular phenotype. DNMT1 is the key enzyme responsible for maintaining methylation patterns during DNA replication. While the role of Dnmt1 has been described in multiple organs, here we identified a novel, critically important mechanism how Dnmt1 controls tissue progenitors. The greatest methylation difference in Dnmt1 knock-out mice was observed on transposable elements (TE), which resulted in increase of endogenous retroviruses and cell death. We believe that release of TE was a critically overlooked component of phenotype development in previous studies that our comprehensive genome wide methylation analysis allowed us to identify.

**Competing interests:** The Susztak lab receives research support from Biogen, Boehringer Ingelheim, Celgene, GSK, Merck, Regeneron and ONO Pharma for work not related to this manuscript.

## Introduction

DNA cytosine methylation (5mC) is largely erased and then re-established between generations in mammals. At the blastocyst stage, due to active and passive demethylation, most cytosines are unmethylated. During cell type diversification and differentiation, CpG methylation reaches about 60% (1). DNA methylation is accomplished by the *de novo* DNA methyltransferases such as DNMT3A and DNMT3B (2, 3). DNMT3A and 3B have been proposed to be critical for establishing methylation of enhancers that are important for cell-type specific transcription factor binding and cell differentiation. DNMT1 is a hemimethylase and therefore it is the key enzyme responsible for maintaining methylation patterns during DNA replication (2, 4). Combination of cytosine methylation, histone modification and chromatin structure define genome accessibility and allows the differentiation of multiple cell types from a single fertilized egg (5).

*De novo* cytosine methylation of regulatory regions (promoters and enhancers) has been proposed to play a key role in establishing gene expression and thereby cellular differentiation and lineage commitment. While this model of gene expression and differentiation is widely accepted, the direct evidence linking promoter/enhancer methylation to gene expression regulation *in vivo* is limited due to technical difficulties in manipulating cytosine methylation in a temporal and site-specific manner. Several studies showed that 5mC is mainly associated with genes that are already repressed by other mechanisms, indicating that 5mC might not be a universal mechanism of repressing gene expression (6, 7). These results indicate that the role of 5mC in regulating gene expression might be more complex than previously appreciated (8).

Animal-model studies with genetic deletion of Dnmts in different cells and organs have been used to understand the role of methylation of regulating gene expression and cellular differentiation. In embryonic stem cells, pancreatic, gut epithelial and skin progenitors, Dnmt1 depletion results in dramatic organ development failure (9-12). The mechanism of Dnmt1 loss-induced organ defect has been attributed to differential methylation of cell type specific genes leading to cell cycle arrest, premature differentiation, and a failure of tissue self-renewal (13, 14). Interestingly, deletion of *de novo* methyltransferases (Dnmt3a or 3b) in different compartments have been associated with milder phenotypes (6).

Transposable elements (TE) control is one of the most ancient and fundamental function of cytosine methylation. TEs are generally fully methylated in all cells (15). Transposons are mobile genetic elements found in every eukaryotic genome sequenced to date and account for at least half of the mammalian genome (16). In most mammals, retrotransposons are the predominant TEs. These can be divided into long terminal repeat (LTR) retrotransposons, including endogenous retroviruses (ERVs), non-LTR retrotransposons such as long interspersed elements (LINEs) and short interspersed elements (SINEs) (15, 17). Due to the low methylation state of the blastomere, some ERVs are highly expressed at this stage. While some believe, that ERV expression is a simple by-product of epigenetic reprogramming, studies indicate that ERVs could contribute to multipotency. Recent studies also found significant ERV expression in some cancer cells due to epigenetic misregulation. Release of TE silencing in cancer cells induces the cytosolic double-stranded RNA (dsRNA) sensing pathway that triggers a Type I interferon response (18, 19).

Nephron number in humans is highly variable, and ranges from 200,000 to 2,000,000. Six2-expressing cap mesenchymal cells represent a multipotent nephron progenitor population, which undergo self-renewal and give rise to the functioning nephron epithelium in the kidney (20). Mammals, unlike some species of fish, are unable to grow or regenerate nephrons and nephron number is set during development. Furthermore, nephron number at birth shows strong association with hypertension and kidney disease development-later in life. Studies have shown that intrauterine nutritional and environmental alterations are the key determinant of nephron endowment and more recent studies indicate that epigenetic changes are the key mediators of developmental programming. Epigenome-modifying enzymes use products of intermediate metabolism (methyl or acetyl groups) as their substrates and variations in substrate availability can lead to changes in the epigenome. Therefore, epigenetic changes during development have been proposed to be the mediators of fetal programming that eventually lead to kidney disease development (21-24). In addition, alterations in histone acetylation caused by loss of Hdac1 and Hdac2 in the ureteric-bud-derived cells result in proliferation and differentiation defect via impaired p53 and Wnt signaling (25, 26).

The current study aimed to understand the role of cytosine methylation in the Six2 positive progenitor population to identify functionally important epigenome-modifying enzymes and regions in the genome where methylation modifications are functionally important for kidney development.

## Results

### Changes in Dnmt and Tet expression in developing kidneys

First, we investigated the temporal course and relative expression of Dnmt and Tet levels in mouse kidney samples. Gene expression analysis (by QRT-PCR) indicated that the expression of Dnmt1, 3a, 3b, Tet1 and Tet3 were higher at birth (P0) compared to adult. We did not observe any temporal change in Tet2 expression (**Figure 1A).** By analyzing RNAsequencing data of P0 and adult mouse kidney samples, we found that the expression of Dnmt3a, Dnmt1, and Tet3 were the highest amongst all epigenome editing enzymes, that were analyzed in P0 mouse kidney samples **(Figure 1B)**.

**Figure 1.**
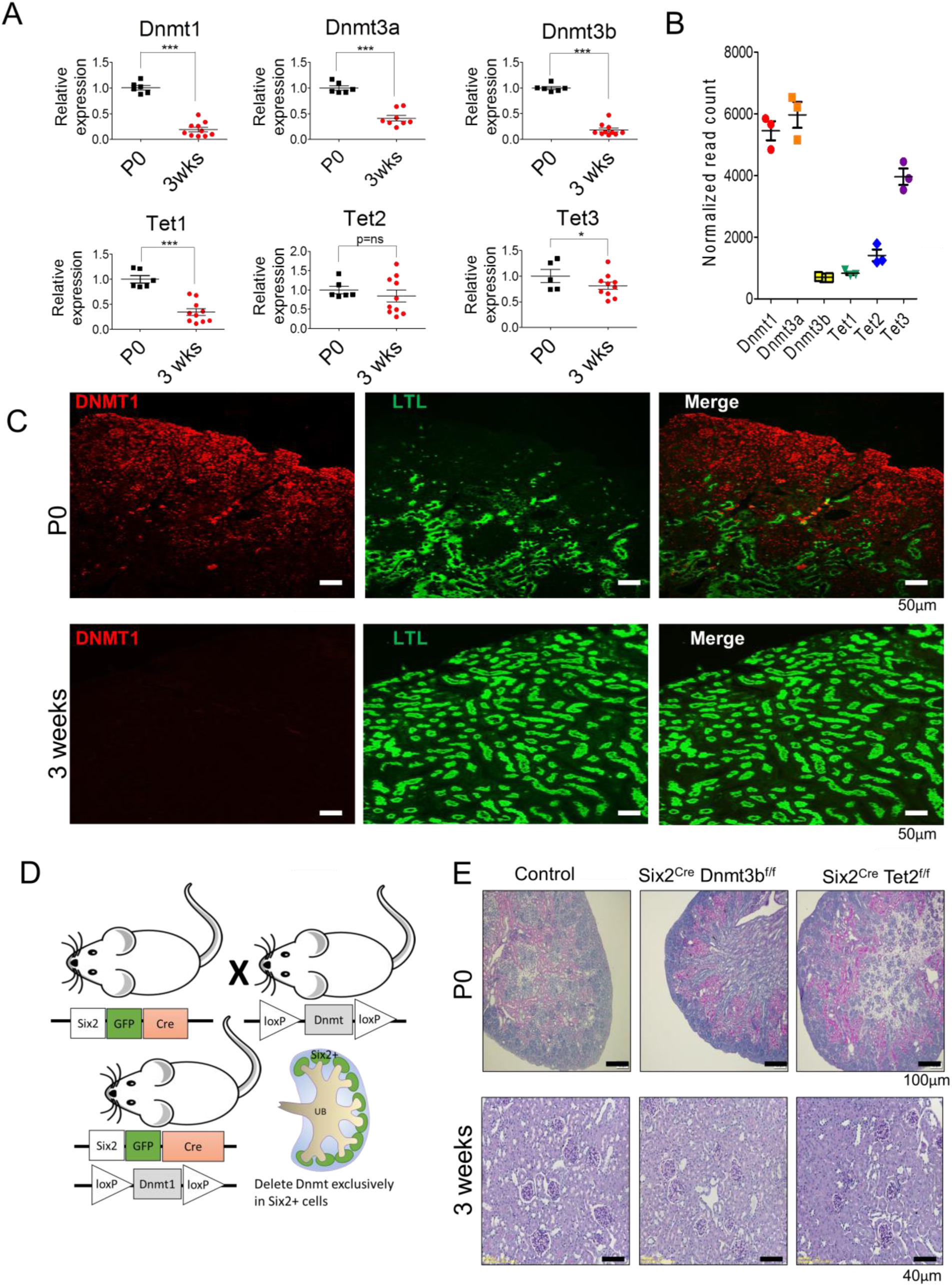
Expression of epigenome editing enzymes in the kidney. Expression of Dnmt1, 3a, 3b, Tet1, 2 and 3 in newborn (p0) and 3 weeks old mouse kidneys analyzed by (**A**) QRT-PCR and (**B**) RNAsequencing. (**C**) Immunofluorescence staining of DNMT1 in P0 and 3 weeks old mouse kidneys. (**D**) Breeding scheme of generating Six2^Cre^ Dnmt1^f/f^ mice. (**E**) Representative PAS stained sections of control, Six2^Cre^ Dnmt3b^f/f^ and Tet2 ^f/f^ kidneys in new born (P0) and 3 weeks old mice.

Protein expression analysis, by immunofluorescence staining, indicated that in the P0 mouse kidney cortex, DNMT1 was mainly expressed in the nephrogenic zone and its expression gradually decreased as epithelial cells differentiated. We failed to detect DNMT1 expression in differentiated proximal tubules (LTL-labeled segments) in developing kidneys. DNMT1 level was also very low in adult kidney samples **(Figure 1C)**. In summary, these results indicate that the expression of methylome editing enzymes decrease during epithelial differentiation in the kidney, as major changes occur mostly during development.

### Dnmt3a, Dnmt3b, Tet1 and Tet2 are dispensable for kidney development

In mice, Six2 positive cells give rise to most of the functional nephron epithelium from glomerular to distal tubule epithelial cells. To determine the functionally significant methylome editing enzymes, we conditionally deleted Dnmt1, 3a, 3b, Tet1 and Tet2 in the nephron progenitor population by crossing mice with a Dnmt1, 3a, 3b, Tet2 conditional alleles with Six2-EGFP^Cre^ animals (20) (**Figure 1D).**

Mice with genetic deletion of Dnmt3a, 3b, and Tet1 and Tet2 (Six2^Cre^/Dnmt3a^flox/flox^, Six2^Cre^/Dnmt3b^flox/flox^, Six2^Cre^/Tet2^flox/flox^ or Tet1-/-) were generated by standard mating. The gene deletion was confirmed by QRT-PCR. These animals showed no clear structural and functional abnormalities even at 16 weeks of age (**Figure 1E)**, indicating that Dnmt3a, 3b, Tet1 and Tet2 are dispensable for kidney development.

### Loss of Dnmt1 in Six2+ cells results in severe kidney developmental defect

To study the role of Dnmt1 in kidney development, we created Six2^Cre^Dnmt1^f/f^ animals. Dnmt1 transcript level in whole kidney lysates of P0 kidneys was about 50% lower in Six2^Cre^Dnmt1^f/f^ kidneys compared to Dnmt1^f/f^ controls (P<0.01) while no change in transcript level of Dnmt3a and 3b was detected **(Figure 2A)**. Immunofluorescence staining of Six2^Cre^Dnmt1^f/f^ samples confirmed the lower expression of DNMT1 in cap mesenchyme cell lineage (podocytes to distal tubules) but not in ureteric bud lineage (collecting ducts in medulla) (**Figure 2B).**

**Figure 2.**
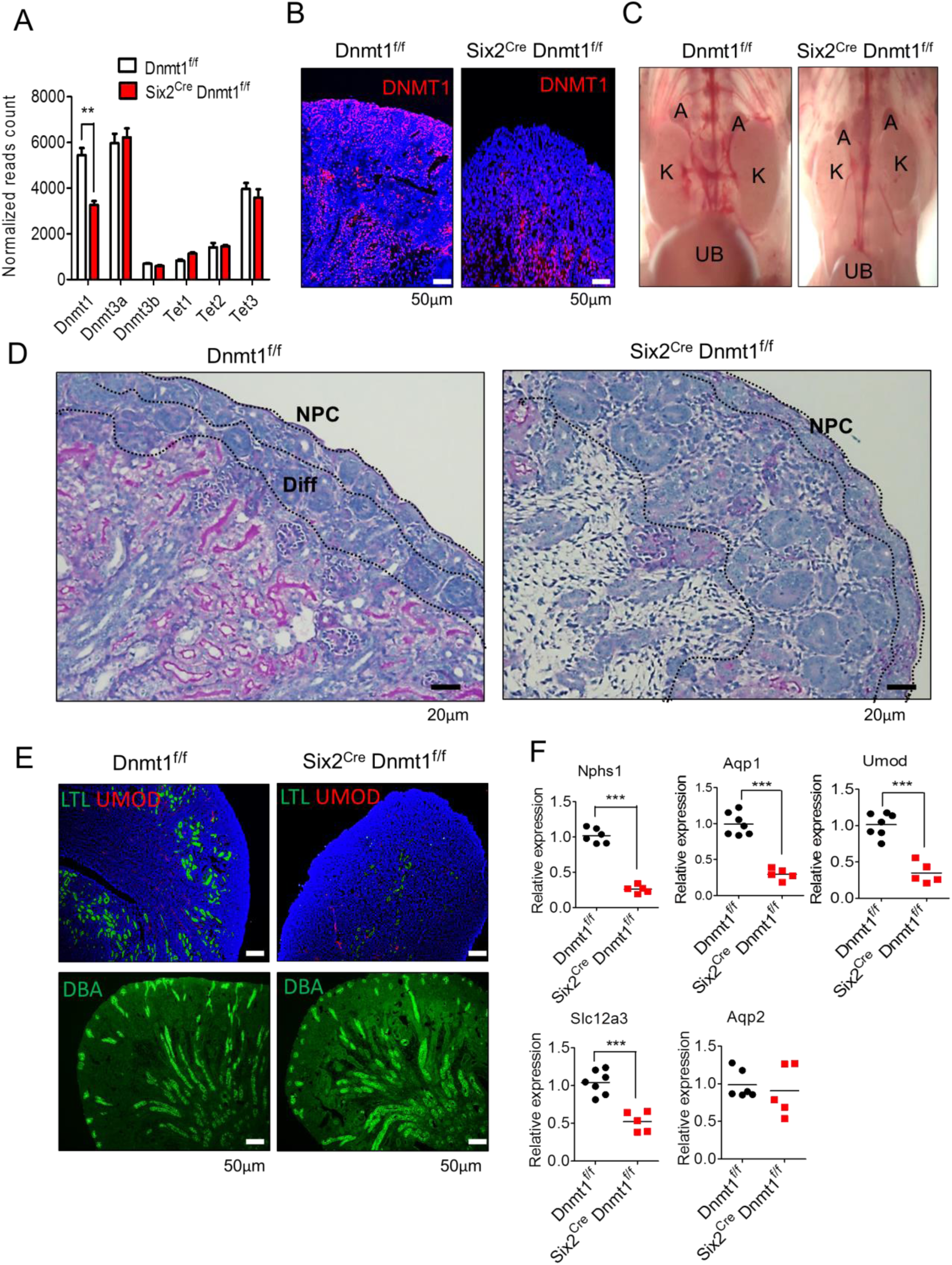
Dnmt1 mediated DNA methylation is critical for kidney development. (**A**) Analysis of RNAsequencing data for the expression of Dnmt1, 3a, 3b and Tet1, 2 and 3 in Dnmt1^f/f^ and Six2^Cre^ Dnmt1^f/f^ mouse kidneys (P0) as analyzed by RNAsequencing. (**B**) Immunofluorescence staining of DNTM1 (red) and nuclei (DAPI, blue) in Dnmt1^f/f^ and Six2^Cre^ Dnmt1^f/f^ kidneys. (**C**) Gross morphology of Dnmt1^f/f^ and Six2^Cre^ Dnmt1^f/f^ mice (urinary bladder: UB, kidney: K and adrenal: A) (**D**) Representative PAS stained kidney sections from control (Dnmt1^f/f^) and Six2^Cre^ Dnmt1^f/f^ kidneys (P0). NPC; nephron progenitors, Diff; differentiation (**E**) Immunofluorescence staining of Dnmt1^f/f^ and Six2^Cre^ Dnmt1^f/f^ kidneys (upper panel LTL (lotus lectin; proximal tubule marker): green, UMOD (uromodulin; Loop of Henle marker): red and DAPI: blue, bottom panel DBA: green). (**F**) Gene expression in Dnmt1^f/f^ and Six2^Cre^ Dnmt1^f/f^ kidneys by QRT-PCR.

Six2^Cre^ Dnmt1^f/f^ mice were born at expected Mendelian ratio, but they died within 24 hours of birth. Animals had visibly smaller kidneys and their bladders were void of urine **(Figure 2C).** Structural analysis indicated significant defect in nephron differentiation. In control P0 mouse kidneys, we were able to observe multiple layers of nascent nephrons as they formed in the nephrogenic zone, followed by a thick layer of renal cortex. Histological analysis of the Six2^Cre^ Dnmt1^f/f^ kidneys, showed some immature renal vesicles, and very few comma and S-shape bodies, but no glomeruli and fully differentiated nephrons **(Figure 2D).**

Immunostaining studies further confirmed the lack of differentiated renal tubule epithelial cells in Six2^Cre^Dnmt1^f/f^ mice. Mature proximal tubule cells stained with LTL in control mice, but there was no LTL-positive segment in kidneys of knock-out animals. UMOD is a marker of differentiated loop of Henle segment. Immunofluorescence studies failed to identify UMOD-positive cells in the Dnmt1 knock-out kidneys. On the other hand, kidney segments differentiated from ureteric bud segments, such as the DBA positive collecting duct cells, appeared to be unaffected (**Figure 2E)**. Gene expression analysis by QRT-PCR showed lower levels of podocyte, proximal tubule, Henle’s loop, and distal loop specific marker gene expression in Six2^Cre^ Dnmt1^f/f^ mice (all p<0.0001) (27). The expression of the collecting duct marker, Aqp2, was unchanged in Six2^Cre^ Dnmt1^f/f^ mouse kidneys **(Figure 2F).** In summary, our studies indicate that no fully differentiated nephrons are formed in the absence of Dnmt1 from Six2+ progenitor cells.

### Dnmt1 depletion did not reduce Six2 positive cell population

To understand the mechanism of Dnmt1 depletion-induced defect in nephrogenesis, we first quantified the number of Six2+ cell population. We used three different approaches. First, we quantified the Six2+ (GFP+) cells in P0 kidneys by fluorescence-activated cell sorting (FACS) as the Cre line also contained the GFP transgene. We found no significant differences in GFP+ cell number in control and Six2^Cre^ Dnmt1^f/f^ mice **(Figure 3A and B).** Second, we performed immunofluorescence staining for SIX2. We observed that Six2^Cre^ Dnmt1^f/f^ mice had slightly, but not significantly higher SIX2+ staining **(Figure 3C).** To understand changes in epithelial differentiation we examined the expression of Six2 and Cited1, genes critical for cell differentiation (28). We found that while Six2 transcript was unchanged, Cited1 expression was decreased in Dnmt1 knock-out kidneys **(Figure 3D and E).** In summary, we conclude that we were unable to identify significant differences in the Six2 expression in the kidney progenitor pool.

**Figure 3.**
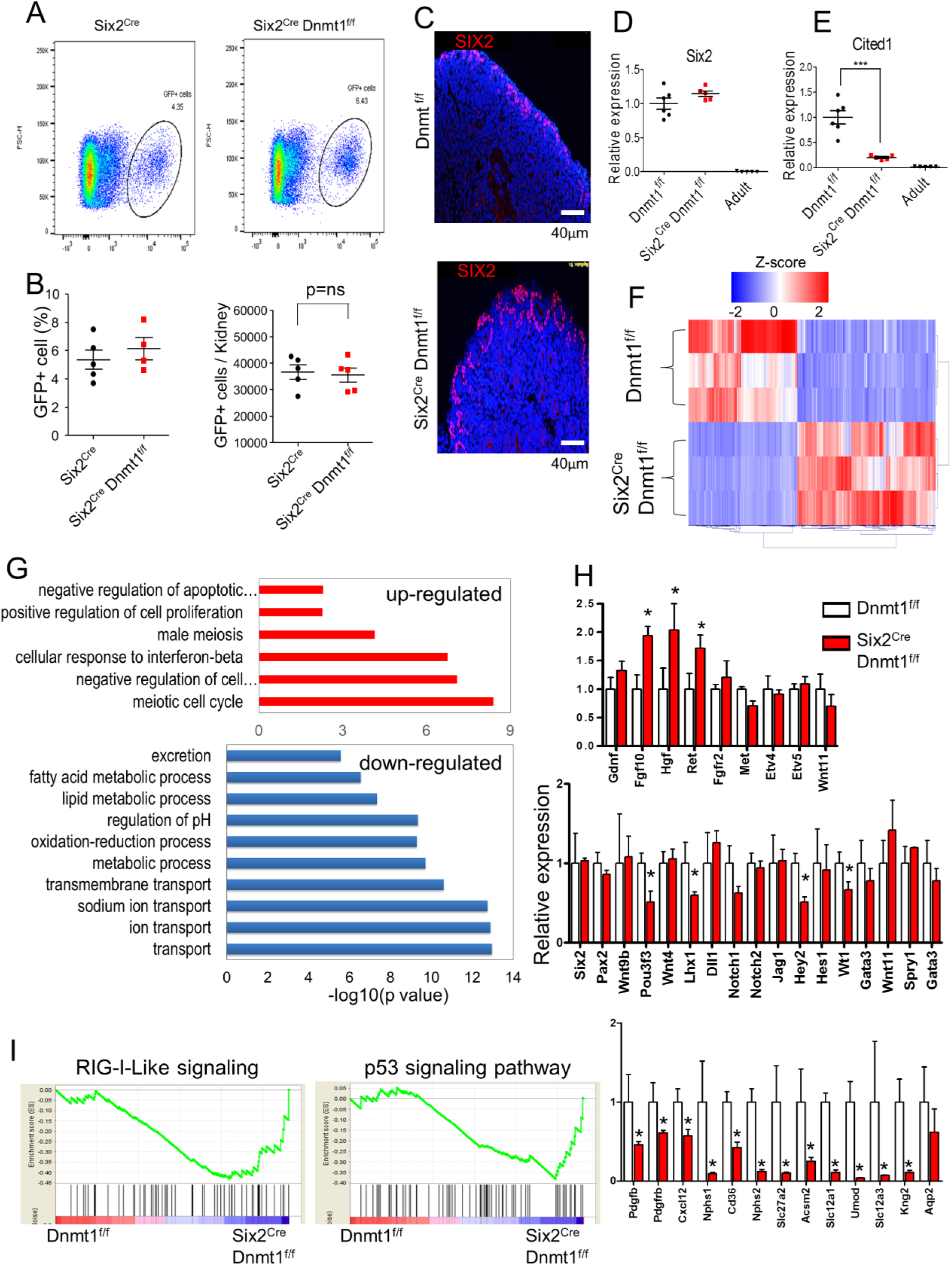
Gene expression analysis of Six2Cre Dnmt1 f/f kidneys. (**A**) Representative flow cytometry images of GFP+ cells in Six2^Cre^ and Six2^Cre^ Dnmt1^fl/fl^ kidneys. (**B**) Quantification of GFP+ cells. (**C**) Immunofluorescence staining of SIX2 in Dnmt1^f/f^ and Six2^Cre^ Dnmt1^fl/fl^ kidneys. (**D and E**) Expression of Six2 and Cited1 in Dnmt1^f/f^ P0, six2^Cre^ Dnmt1^f/f^ P0 and wild type adult kidneys, as analyzed by QRT-PCR. (**F**) Heatmap of the differentially expressed genes between Dnmt1^fl/fl^ and Six2^Cre^ Dnmt1^fl/fl^ mice. Color scheme is based on z-score distribution. (**G**) Functional annotation analysis (DAVID) of the differentially expressed (up- and down-regulated) genes. (**H**) Relative expression level of genes involved in kidney development. Y-axis is normalized expression level determined by RNA sequencing. Asterisks represent significant differences calculated by DESeq2. (**I**) The GSEA enrichment plots of RNAsequencing data showing the distribution of genes in the sets that are increased in Six2^Cre^ Dnmt1^fl/fl^ mice.

In order to understand the role of Dnmt1 in kidney development, we performed comprehensive gene expression analysis (by RNAsequencing). We identified 1,829 differentially expressed genes (FDR<0.05) when we compared P0 kidneys of Dnmt1 knock-out mice to controls **(Figure 3F; Table S1)**. Genes that showed significantly lower expression in knock-out kidney samples were related to ion transport and renin-angiotensin pathways **(Figure 3G; Figure S1),** indicating the lack of renal tubule epithelia in the knock-out mice. While Six2 expression was unaffected, we found that the level of Hgf, Fgf10 and Ret were increased in Dnmt1 null kidneys (**Figure 3H**). These genes are usually expressed prior to Six2, as they are critically involved in the ureteric bud and metanephric mesenchyme interaction and the final establishment of Six2 positive cap mesenchyme nephron progenitors. Levels of genes that involved in mesenchymal to epithelial transition such as Wnt4 and Pax2 and others did not all show significant differences (**Figure 3H**). Genes that have been shown to play role in elongation and segmentation of the renal epithelial vesicles or markers of differentiated glomerular and proximal tubule cells were almost uniformly decreased in Dnmt1 knock-out mice (**Figure 3H**). Transcription factors that maintain collecting duct cell fate, such as Aqp2 were not changed in Six2^Cre^ Dnmt1^f/f^ mice, as this segment develops from a Hoxb7 and not from a Six2 positive population. These results indicate an absence of differentiation of Six2 progenitor cells and re-expression of early nephric duct markers.

We found that male primordial germ cell specific genes, such as Dazl and Sohlh2 showed one of the highest increases in Dnmt1 null kidneys (**Figure S2A and B**). There was an enrichment for genes with function in homologous recombination and meiosis in absence of Dnmt1 (**Figure 3G; Figure S1**). We observed the expression of even earlier markers such as Zscan4b and Zscan4d that are normally only expressed at 2-4 cell-stage embryo (29). The full list of differentially expressed genes can be found under **Table S1**. Gene Set Enrichment Analysis (GSEA) was performed using RNAsequencing data to identify the signaling pathways changed in absence of Dnmt1. An increase in expression of interferon and retinoid acid inducible gene I (RIG-I) signaling, which are viral pattern recognition receptors, was observed in kidneys in absence of Dnmt1. Increase in expression p53 and associated cell death has been previously described in absence of Dnmt1 and a robust difference in p53 pathway was also evident in kidney samples (**Figure 3I; Figure S1 and S3**). In summary, genome-wide expression analysis not only indicate the lack of differentiated epithelial cell markers but also the expression of early 2-4 cell-stage embryo markers and viral pattern recognition genes in absence of Dnmt1.

### Genome wide methylation during kidney development

In order to understand the role of changes in 5mC in kidney development, we have first analyzed global methylation changes in newborn and adult kidney tissue samples. As global methylation analysis has not been reported in developing mouse kidneys, we used reduced representation of bisulfate sequencing (RRBS) to enrich the methylation analysis of regulatory regions. 5mC levels were slightly but significantly lower in adult kidney samples when compared to newborn kidneys (43.7% adult vs 50.5% fetal p=0.0032) that are still undergoing nephrogenesis. We have identified 8,345 differentially methylated regions; DMR (4,178 with higher while 4,167 with lower 5mC in adult, FDR<0.05) **(Figure 4A)**. Upon mapping cytosine methylations to RefSeq-based regulatory annotation we found low 5mC levels around transcription start sites (TSS) (**Figure 4B**).

**Figure 4.**
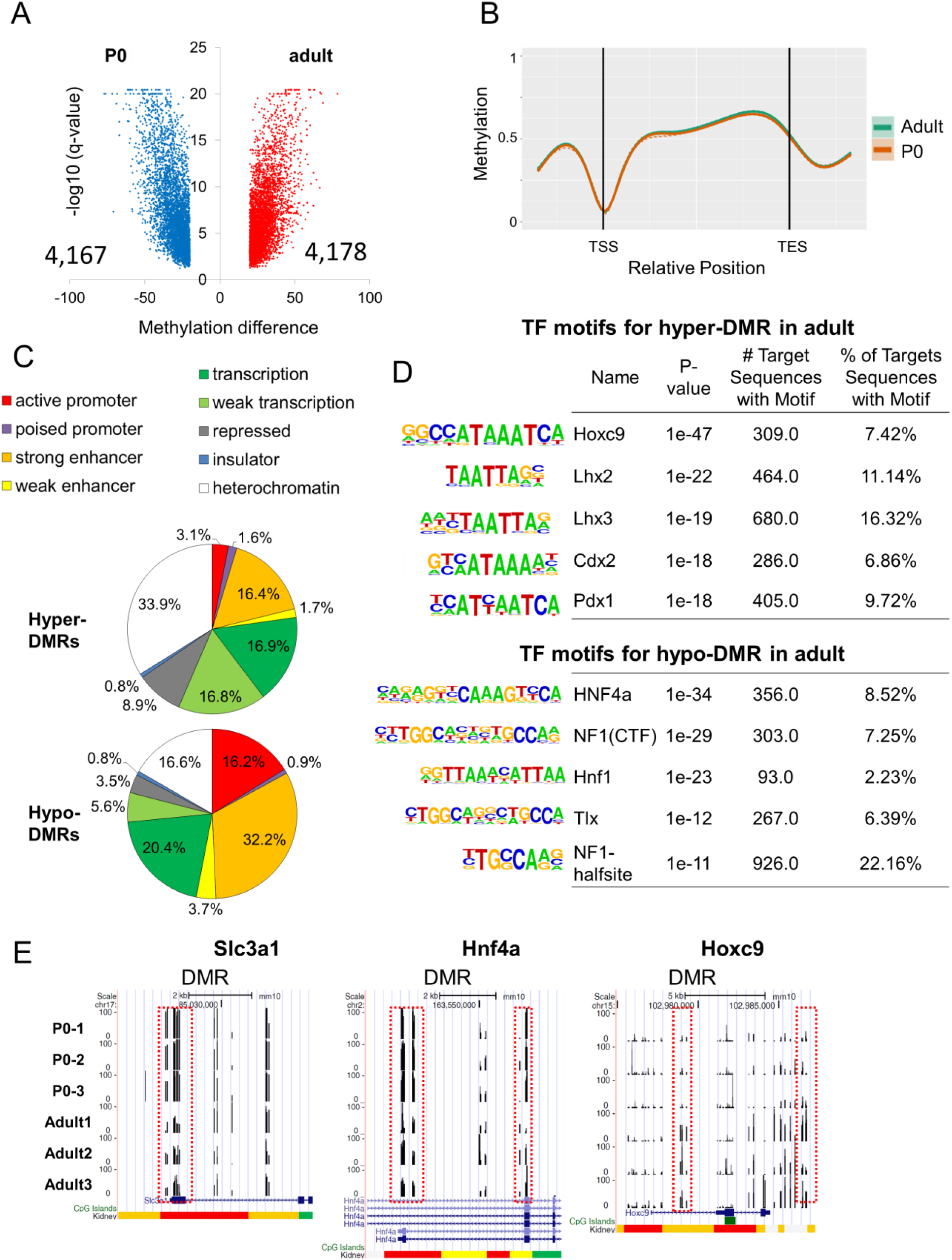
Genome wide methylation changes during kidney development. (**A**) Volcano plot showing the differentially methylated regions (DMRs) between kidneys of control P0 and adult mice. (**B**) Methylation profiles near and within all mouse genes. TSS: transcription start site. TES: transcription end site. (**C**) Annotation of DMRs by kidney-specific chromatin states. (**D**) Transcription factor motif enrichment analysis of DMRs. (**E**) UCSC genome browser screenshot showing the representative DMRs. DMRs are marked by red boxes. Y-axis represents methylation level from 0% to 100%. Chromatin states are shown at the bottom.

Next, we mapped DMRs onto kidney specific functional regulatory regions, defined by genome-wide histone tail modification mapping (ChIP-Seq) and computational integration of multiple marks using ChromHMM (30, 31). Promoters and enhancers were significantly enriched in regions showing lower methylation levels in adult kidney samples **(Figure 4C**), whereas heterochromatin regions were the most significantly enriched groups that showed an increase in 5mC levels in adult kidney samples. We further identified key transcription factor binding motifs in the DMRs. Regions with increase in their methylation levels were enriched for developmental transcription factors, such as Lhx binding sites. On the other hand, methylation levels were lower of sites where Hnf was binding in adult kidneys, as Hnf is one of the key (cell identity) transcription factor for differentiated renal tubule epithelial cells **(Figure 4D).** Gene ontology analysis for DMRs with increased methylation, indicated enrichment around pathways involved in embryonic morphogenesis and development **(Figure S4A),** and decreased in development and cell shape control (**Figure S4B**). For instance, DMRs were detected in Slc3a1, Hnf4a and Hoxc9 gene loci **(Figure 4E)**. In summary, our results indicate significant methylation changes of regulatory regions (promoter and enhancer) regions in the developing kidney.

### Dnmt1 loss results in broad cytosine methylation changes

Having established that Dnmt1 is required for normal kidney development, we analyzed DNA methylomes of control and knockout kidneys. Pair-wise comparison of base resolution, genome-wide reduced representation bisulfate sequencing (RRBS) CpG methylation plots revealed similar CpG methylation profiles within control or knockout groups. Cytosine methylation showed a bimodal distribution in control samples. There was a marked decrease in the number of fully methylated loci in knock-out kidney samples (**Figure 5A).** We observed a significant decrease in overall methylation (50% in control and 37% in absence of Dnmt1 p=0.004) (**Figure 5B)** and most of DMRs were hypo-DMRs (45,026 hypo vs 88 hyper). Next, we mapped DMRs onto kidney specific regulatory regions, defined by histone modifications and ChromHMM maps. As illustrated in **Figure 5C**, surprisingly only a few promoters or enhancers showed differential methylation in absence of Dnmt1. Identification of DMR for transcription factor motifs revealed enrichment for Elk4, Mef2c and Gata4 binding motifs (**Figure 5D**); these are cardiac and myocyte specific transcription factors that are normally not expressed in the kidney. Lower methylated regions in Dnmt1 knock-out mice were enriched around genes with action potential and filament depolymerization functions (**Figure S4C**), showing consistency with the TF binding data.

**Figure 5.**
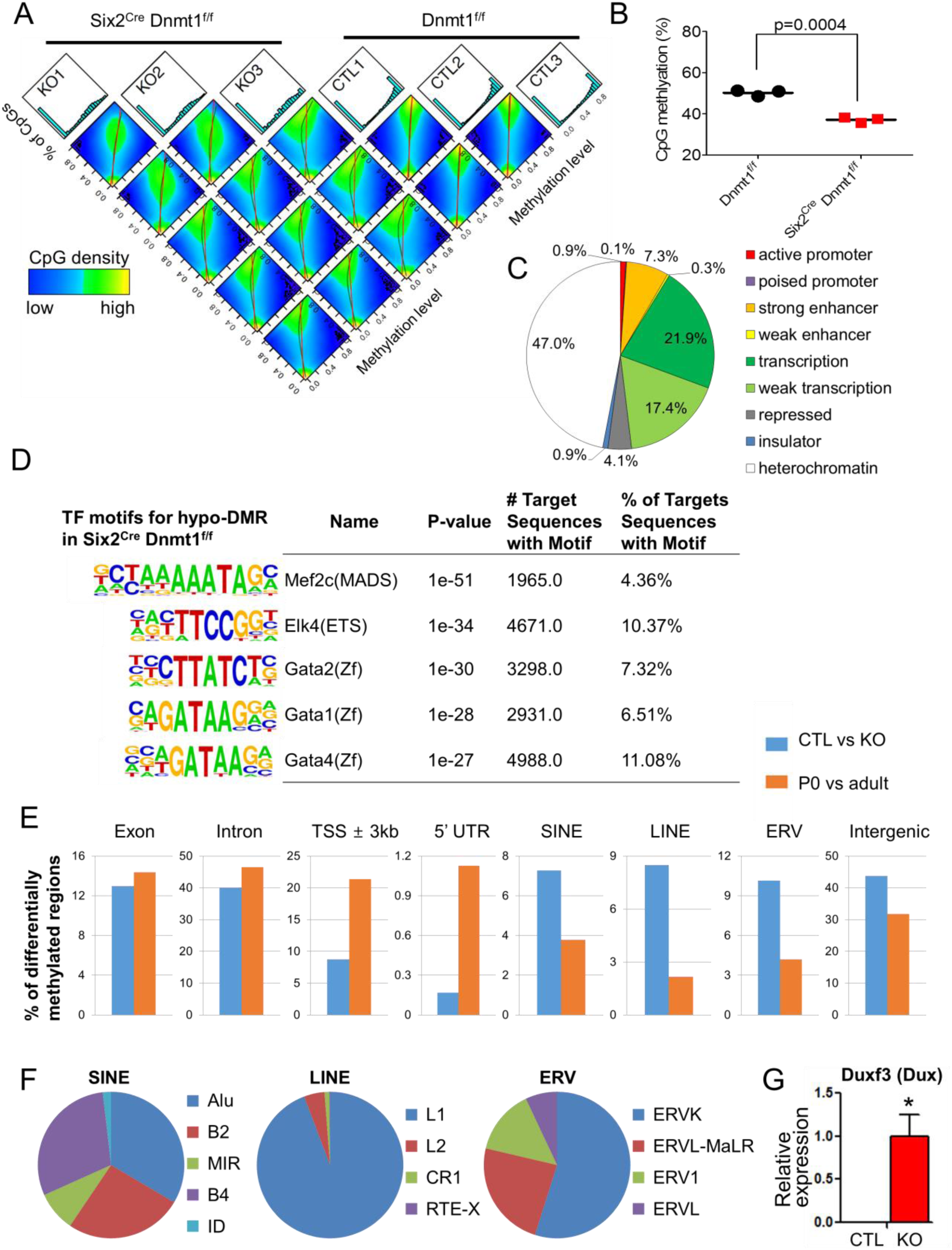
Genome wide methylation changes in Six2^Cre^ Dnmt1fl/fl kidneys. (**A**) Pairwise correlation analysis of Dnmt1^fl/fl^ and Six2^Cre^ Dnmt1^fl/fl^ mouse kidneys. Histogram plots showing counts by CpG methylation level (x axis: 0% to 100%) distributed across 20 bins of 5% intervals. Scatter plots showing correlation between every pair of samples. (**B**) Global CpG methylation in Dnmt1^fl/fl^ and Six2^Cre^ Dnmt1^fl/fl^ mouse kidneys. (**C**) Annotation of DMRs by kidney-specific chromatin states. (**D**) Transcription factor motif enrichment analysis of DMRs showing lower methylation levels (hypo DMR). (**E**) Distribution of DMRs in different genomic elements. Blue bar represents DMRs detected between Dnmt1^fl/fl^ and Six2^Cre^ Dnmt1^fl/fl^ mice. Red bar represents DMRs detected between Dnmt1^fl/fl^ P0 and wild type adult mice. **(F)** Distribution of DMRs in different transposable elements. **(G)** Relative expression level of Duxf3 (Dux). Y-axis is normalized expression level determined by RNA sequencing. Asterisks represent significant differences calculated by DESeq2.

More than 50% of the observed DMRs in the Dnmt1 knock-out mice were on regions annotated as repressed region or heterochromatin (**Figure 5C**). In absence of Dnmt1, methylation differences of intergenic regions, including LINE, SINE and ERV were the most prominent **(Figure 5E).** LTR retrotransposons including endogenous retroviruses (ERVs) derived from infectious retroviruses that integrated in the germline. Most of DMRs detected in ERV were hypo-DMRs (4,577/4,579) and more than half of them were observed in ERVK family (**Figure 5F**). In contrast, transcription start sites and 5’ UTR showed differential methylation during development (**Figure 5E**). Furthermore, in Dnmt1 knock-out mice we detected significant increase of expression level of Dux transcription factor which is known as a ERV elements activator (**Figure 5G**) (32, 33). In summary, loss of Dnmt1 was associated with significantly lower methylation levels in the kidney and resulted in significant loss of methylation in heterochromatin regions (LINE, SINE and ERV), while significant differences were observed in methylation of promoters and enhancers during normal development.

### Dnmt1 maintains silencing of transposable elements in Six2+ progenitor cells

Next, we integrated methylation changes with transcriptome differences. In Six2^Cre^ Dnmt1^f/f^ kidneys, 45,114 DMRs and 1,829 differentially expressed genes were identified (**Figure 6a**). Large numbers of differentially expressed genes (749 of the 1,829) were in the vicinity of DMRs. Number of genes (n=415) showed decrease both in methylation and gene expression levels. Genes in this group, mostly included renal tubule epithelial specific genes such as Scl and Umod, consistent with the loss of differentiated epithelial cells in Six2^Cre^ Dnmt1^f/f^ kidneys (**Figure 6B; Figure S2A**). Expression of these genes were lower despite the low methylation in their regulatory regions, indicating lack of clear correlation between methylation and gene expression. Most of such genes, were markers of differentiated renal epithelial cells and transporters **(Figure S5A)**. A small fraction (18%) of all differentially expressed genes (334) followed a classic pattern of lower methylation and increase in gene expression. Gene ontology analysis showed this cluster of genes are enriched for meiotic cell cycle, and spermatogenesis. **(Figure S5B)**. Dnmt1 KO kidneys expressed high levels of primordial germ cell (PGC)-specific genes including Dazl (34, 35), Sohlh2 (36, 37), Tex19.1 (38), Pet2 (39) and also genes exclusively expressed in 2-4 cell-stage embryo, Zscan4b and Zscan4d (29). Such genes were undetectable in control P0 kidneys **(Figure 6B; Figure S2A).** Tissue enrichment analysis of the differentially expressed genes indicated enrichment for brain, testis and embryo genes **(Figure 6C),** highlighting a broad dysregulation in cell identity in Dnmt1 null kidneys.

**Figure 6.**
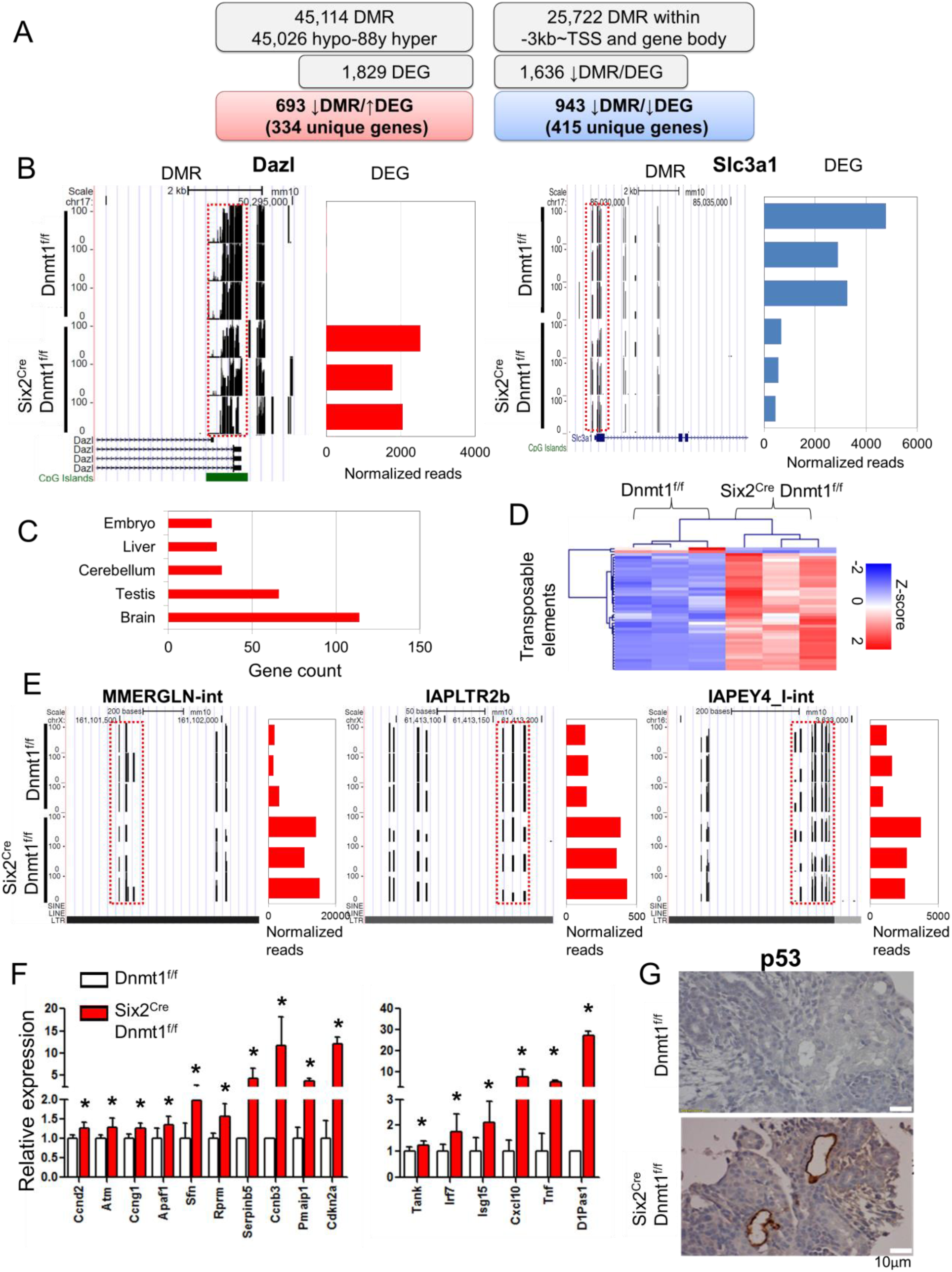
Correlation between methylation and gene expression changes in Dnmt1 null kidneys. (**A**) The number of identified differentially methylated region (DMR) and differentially expressed genes (DEG) and their consistency and directionality in control and Six2^Cre^ Dnmt1^fl/fl^ mice. (**B**) Representative DMRs and their target gene expression changes. DMRs are marked by red boxes. Left panel shows a UCSC genome browser screenshot with DNA methylation patterns and Y-axis represents methylation level from 0% to 100%. Right panel shows gene expression level and X-axis represents normalized sequencing reads. (**C**) Tissue enrichment analysis of the target genes (DAVID). (**D**) Heatmap of the differentially expressed transposable elements (TEs) between Dnmt1^fl/fl^ and Six2^Cre^ Dnmt1^fl/fl^ mice. Color scheme is based on z-score distribution. (**E**) Representative DMRs and their target TE expression changes. DMRs are marked by red boxes. Left panel shows a UCSC genome browser screenshot with DNA methylation patterns and Y-axis represents methylation level from 0% to 100%. Right panel shows TE expression level and X-axis represents normalized sequencing reads. (**F**) Relative expression level of genes involved in RIG-I-like and p53 signaling. Y-axis is normalized expression level determined by RNA sequencing. Asterisks represent significant differences calculated by DESeq2. (**G**) P53 immunohistochemistry staining of kidneys of control (Dnmt1^fl/fl^) and Six2^Cre^ Dnmt1^fl/fl^ mice.

Given that methylation changes were enriched on TE in the Dnmt1 null mice, we have re-aligned the mouse RNAsequencing data (see Materials and Methods for detail) so we can understand whether reduced methylation on transposable elements were associated with increased ERV transcription. We found that many ERV and 3 LINE transcript levels were higher in Dnmt1 knock-out kidneys. Among 619 LINE, SINE and ERV families analyzed we have identified 43 differentially expressed transposable elements, of which 38 ERV families were increased and only 2 ERV families were decreased in Dnmt1 KO samples **(Figure 6D and Table S2)**. These increased ERV families included all 3 ERV classes. On the other hand, 3 LINE families (L1Md_Gf, L1Md_T and L1Md_F) were increased in Dnmt1 KO samples. These LINE families were relatively young and most had an F-type promoter (40). **Figure 6E** illustrates Dnmt1 depletion-induced hypomethylation was associated with increase in RNA levels of 3 different endogenous retroviruses, MMERGLN, IAPLTR2b and IAPEY4.

While pluripotent stem cells have been shown to have an increase in ERV levels that might even be functionally important during development and differentiation, increase in ERV in cancer cells triggers cytosolic viral sensors and interferon release and immune activation. We found that in the developing kidneys increase in ERV levels is associated with increased interferon and RIG-I signaling **(Figure 6F)**. RIG-I-like receptor is a dsRNA helicase that functions as a pattern recognition receptor. The increased in expression of p53 pathway observed in Dnmt1 null mice (**Figure 6F and G**) is likely a key contributor to cell death and subsequent abnormal kidney development in the Dnmt1 null mice.

### Dnmt and Tet are dispensable in differentiated glomerular epithelial cells

To study the role of 5mC in differentiated podocytes, we crossed Dnmt1, 3a, 3b and Tet2 ^loxP/loxP^ animals with the Pod^cre^ mice (41) to generate podocyte specific knockout animals. Mice were born at expected Mendelian frequency and we did not observe significant functional or structural renal abnormalities even at 20 weeks of age **(Figure 7A)**, suggesting these enzymes are not essential for maintaining podocyte functions.

**Figure 7.**
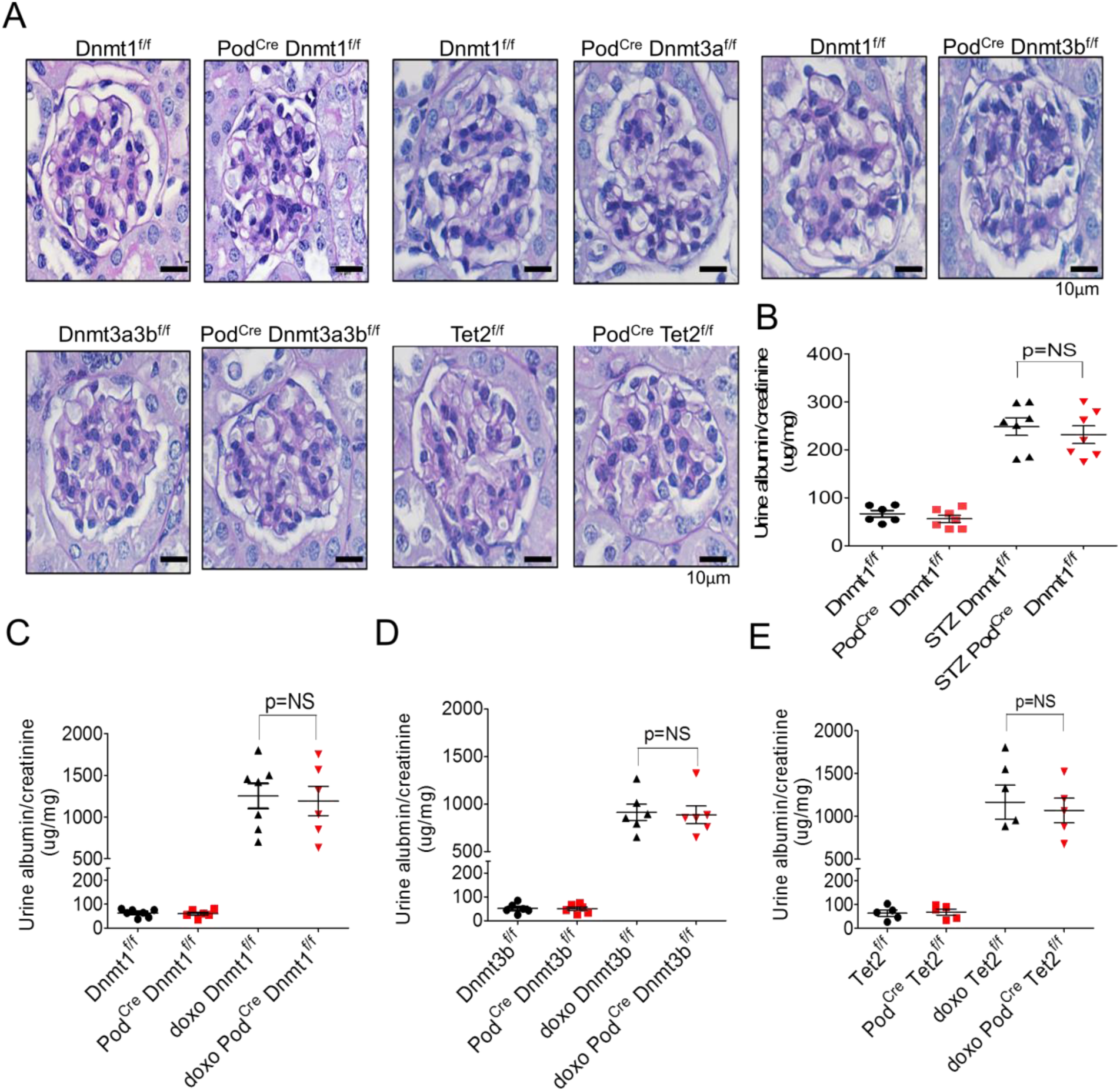
Epigenetic editing enzymes are dispensable in mature podocytes. (**A**) Representative PAS-stained mouse kidney sections of control (Dnmt1^f/f^) and mice with podocyte-specific deletion of Dnmt1, Dnmt3a, Dnmt3b, Dnmt3a/3b and Tet. (**B**) Albuminuria (urine albumin/creatinine ug/mg) of Dnmt1^fl/fl^ and Pod^Cre^ Dnmt1^fl/fl^ mice at baseline and 20weeks following STZ-induced diabetes. (**C-E**) Albuminuria (urine albumin/creatinine ug/mg) of control, Pod^Cre^ Dnmt1^fl/fl^, Pod^Cre^ Dnmt3b^fl/fl^ and Pod^Cre^ Tet2^fl/fl^ mice at baseline and 7 days following doxorubicin-induced glomerular injury.

Next, we tested whether podocyte-specific deletion of Dnmts and Tets would alter the cell’s injury response. We injected podocyte-specific knock-out mice and control littermates with streptozotocin to induce diabetes. Phenotype analysis was performed twenty-four weeks after the initiation of diabetes. We observed no differences in albuminuria levels or structural damage such as mesangial expansion or glomerulosclerosis **(Figure 7B)**.

Doxorubicin (Adriamycin) is known to be toxic to podocytes and can induce focal segmental glomerulosclerosis (FSGS). We compared the injury severity between podocyte–specific Dnmt1, 3b and Tet2 null and control animals in doxorubicin-induced FSGS model. Even after careful characterization of control and knock-out animals, we failed to observe differences in functional parameters such as albuminuria or structural changes such as mesangial expansion or glomerulosclerosis in podocyte specific knock-out animals and their littermate controls **(Figure 7C-E)**. These finding indicated that Dnmt1, 3b and Tet2 proteins are dispensable in podocytes, even under injury condition.

## Discussion

Overall, this is the first study to define the methylome at base-pair resolution in the developing and adult mouse kidneys. To define functionally critical methylation changes we have complemented the methylome atlas by deleting most epigenome editing enzymes. Our current study shows that Dnmt1 plays a critical, non-redundant role in the Six2 positive kidney progenitor population. Dnmt1 loss was associated with marked decrease in methylation levels. We did not observe phenotypic changes in absence of active demethylating enzymes such as Tet1 and Tet2 or upon deletion of Dnmt1 in post-mitotic cells such as podocytes. As Dnmt1 is a hemi-methylase, the global loss of cytosine methylation in Six2^Cre^ Dnmt1^f/f^ was likely the consequence of dilution by cell division, indicating that passive methylation is the major determinant of methylation maintenance.

It was to our surprise that genetic deletion of Dnmt3a or 3b in Six2 positive cells did not result in phenotype development. It is possible (albeit not supported by our results) that Dnmt3a and 3b can compensate for one another. Cell type specific or pioneering transcription factors play greater role in establishing the cell type specific transcriptome of cellular differentiation. While further studies needed to determine base resolution methylation levels in Dnmt3a and 3b animals, these results indicate smaller contribution of *de novo* regulatory region methylation to gene expression and cell type diversification.

Differential methylation of promoters and enhancers showed the most pronounces changes during kidney development, consistent with the well accepted model of cell differentiation. On the other hand, we observed little differential methylation of promoters and enhancers in the Dnmt1 knock-out mice. We observed two broad classes of, most likely related, events. While cells did not lose the expression of Six2, our data showed that in the absence of Dnmt1, Six2+ linage cells regress to early progenitor cells. Gene ontology analysis of Dnmt1 KO kidneys indicated enrichment in brain, testes, and early embryonic genes. There was a striking enrichment for genes with function in homologues recombination and meiosis (GC-specific genes). Given the common lineage between the renal and genital systems, methylation seems to be critical to maintain of stem cell differentiation. On histological examination, the nephron progenitor layer is indistinguishable between control and Six2^Cre^Dnmt1^f/f^ kidneys, however, Dnmt1 null cells formed blastocyst-like structures instead of differentiated nephron segment and increase in expression of the very early metanephric mesenchyme markers. The phenotype development is the likely consequence of such broad changes rather differences in a single gene or even a single transcription factor.

In addition, our data suggest that the most profound effect on phenotype development was caused by lowering the methylation of transposable element. Methylation appears to be essential for TE silencing and the lower methylation was associated in an increase in transcriptional activity of these regions, such as ERV expression, which have been also reported in early embryonic development (14, 42, 43). While ERV activation is important in pluripotent stem cells (44), it seems that Six2+ progenitor cells are unable to tolerate cellular dsRNA. The presence of cellular dsRNA induces an interferon response in cells (45), such as observed in Six2^Cre^ Dnmt1^fl/flx^ mice. In our review of the literature, this is the first study describing ERV activation precipitated by loss of Dnmt1 in tissue progenitor cells. Release of transposable elements has recently been identified as major mechanism of anti-tumor effect of azacitidine in various cancer models (18, 19). Ultimately, ERV activation is associated with apoptosis, such as Trp53 activation in Six2 positive renal progenitor cells. These downstream pathways are consistent with previous descriptions in other organ/cell type systems (46). Haploinsufficiency of p53 has partially protected from phenotype development in Dnmt1 depleted pancreas (11).

Lack of phenotype development following Dnmt1, Dnmt3a, 3b, Tet1 and Tet2 deletion in podocytes at baseline and after injury was again surprising. It is possible that some of these enzymes have overlapping functions, hence future studies using double knock-out animals will be essential to exclude this possibility. On the other hand, our studies indicate minimal remodeling of the podocyte methylome at baseline and disease condition. Therefore, our results do not support the notion that targeting the podocyte methylation could be an important therapeutic strategy for kidney disease prevention.

In summary, this study is the first comprehensive methylation analysis during kidney development. We show the expression of DNA methylation editing enzymes and cytosine methylation profiles changes during nephron maturation. Dnmt1 plays a key role in kidney development but mostly not via a modifying the methylome of cell differentiation genes. Cell division mediated methylation dilution appears to be the greatest determinant of the methylome and phenotype development. Absence of Dnmt1 results in release of transposable element methylation, expression of ERVs and other early progenitor specific genes. The tissue specific progenitor population is unable to tolerate ERV reactivation and induces immune activation and cell death. Overall changes in Dnmt1 during development could play important role in nephron endowment and hypertension and kidney disease development.

## Materials and Methods

### Mice

Dnmt1^flox^ mice were obtained from Mutant Mouse Regional Resource Center (MMRRC_014114-UCD). Dnmt3a^flox^ and Dnmt3b^flox^ mice were obtained from JAX lab (47, 48) Tet2^flox^ mice were purchased from JAX lab (stock number 017573). Tet1 null mice were obtained from JAX lab (stock number 017358) Dnmts and Tet2 mice were crossed to transgenic mice carrying the Pod^Cre^ (14) (JAX stock number 008205) or Six2^Cre^ (JAX stock number 009606). Double transgenic mice and animal gender were identified by genomic PCR analysis (49). Podocyte injury was induced in 6-week-old male mice by doxorubicin (Adriamycin, Pfizer, 20mg/kg, intravenous) or streptozotocin injection (Sigma-Aldrich, 50 mg/kg intraperitoneal, for 5 days).

### Histological procedures and staining

Renal histological changes were examined by PAS stained sections. Immunostaining were performed using the following primary antibodies DNMT1 (sc-271729, Santa Cruz), SIX2 (11562-1-AP, Proteintech), p-53 (sc-6243, Santa Cruz), UMOD (SAB1400296 Sigma), fluorescein labeled lotus tetragonolobus lectin (LTL) and Dolichos Biflorus Agglutinin (DBA) (FL-1321, FL-1031 Vector) after heat-induced antigen retrieval by Tris-EDTA buffer (pH 9.0). Sudan Black was used to block auto-fluorescence.

### Isolation of Six2+ cells

Kidneys from Six2^Cre^ Control and Six2^Cre^ Dnmt1^f/f^ mice were dissected under stereo microscope and placed in RPMI, briefly dissociated using 18G and 21G needles. Kidneys were incubated in collagenase for 10 min at 37C. After being passed though 40um cell strainer, 10% serum was added to neutralize collagenase. After red blood cell lysis, single cell suspension was collected by centrifugation and suspended in 500 ul of PBS containing 2% serum. FACS analysis was performed using BD FACSAria II.

### Reduced representation bisulfite sequencing

Genome wide DNA methylome analysis was performed of using the Premium Reduced Representation Bisulfite Sequencing (RRBS) Kit (Diagenode) following manufacturer’s instruction. In brief, genomic DNA was extracted from whole kidneys using Puregene Core Kit A (Qiagen), 100ng genomic DNA was then digested using the methylation-insensitive enzyme MspI, fragment ends were filled in and adapters were ligated. After size selection by AMPure XP beads, fragments were subjected for bisulfite conversion. Libraries were sequenced on Illumina Hiseq 50bp SE configuration. The adapter sequences and low-quality base calls (< 30) were trimmed from the 3’ end using Trim_Galore program with –rrbs option. The bisulfite converted sequencing reads were then aligned to the mouse reference genome (mm10) using Bismark program (default parameters). Differentially methylated regions (DMRs) was determined using methylKit in R package with definition number of CpG ≥3 and DMC ≥1 and q <0.05. Chromatin states of DMRs were annotated using mouse kidney specific ChromHMM maps (30, 31). Transcription factor motif were identified in the DMRs using Homer program.

### RNA sequencing

RNA was isolated using RNeasy mini kit (Qiagen). 200ng total RNA was used to isolate poly A purified mRNA using the Illumina TruSeq RNA Preparation Kit. Sequencing was performed in Illumina by paired-end 150bp, and the annotated RNA counts (fastq) were calculated by Illumina’s CASAVA 1.8.2. Sequence quality was surveyed with FastQC. Adaptor and lower-quality bases were trimmed with Trim-galore. Reads were aligned to the Gencode mouse genome (GRCm38) using STAR-2.4.1d. The aligned reads were mapped to the genes (GRCm38, version 7 Ensembl 82) using HTSeq-0.6.1. DESeq2 was used to test for differential gene expression between control and transgenic groups.

To analyze the transposable elements, the sequencing reads aligned to the mouse genes were firstly removed. First, the aligned reads were mapped to the genes (GRCm38, version 7 Ensembl 82) with intersection-nonempty parameter for the mode option. Then, the reads which were mapped to the Ensembl genes (marked as_no_feature) were filtered out from the sam files. Finally, the rest of the sequencing reads were aligned to the Repeatmasker which was downloaded from the UCSC genome browser. In regarding to QRT-PCR, 1 µg RNA was reverse transcribed using the cDNA archival kit (Life Technology), and QRT-PCR was run in the ViiA7 System (Life Technology) machine using SYBR Green Master Mix and gene-specific primers **(Figure S6)**. The data was normalized and analyzed using the ΔΔCt method.

### Statistical analysis

Statistical analyses were performed using GraphPad Prism software (GraphPad Software Inc., La Jolla, CA). All values are expressed as mean and standard deviation. Two tailed student t-test was used to compare 2 groups. One-way ANOVA with post-hoc Turkey test was used to compare multiple groups. A p-value less than 0.05 was considered statistically significant.

### Study approval

All experiments in animals were reviewed and approved by the Institutional Animal Care and Use Committee of University of Pennsylvania and were performed in accordance with the institutional guidelines.

## Acknowledgement

**Funding:** Li SY received research support from Taipei Veterans General Hospital –National Yang-Ming University Excellent Physician Scientists Cultivation Program 104-V-A-003 and Ministry of Science and Technology R.O.C MOST 104-2314-B-075 -042 -MY3 and 105-2314-B-075 -050 -MY3. Dr. Park is supported by American Diabetes Association Training grant #1-17-PDF-036. Work in the SusztakLab is supported by the NIH R01 DK076077, DK087635 and DP3 DK108220.

**Author contribution:** SYL and KC created transgenic mouse models, SYL performed animal experiments. JP analyzed the RRBS and RNAseq data. RS performed FACS analysis and immunostaining. MBP performed histopathological analysis. SYL and KS conceptualized the work and wrote the manuscript. SYL and JP contributed to this paper equally.

## Data access

All sequencing data (RRBS and RNAsequencing) have been deposited in NCBI’s Gene Expression Omnibus and are accessible through GEO Series accession number GSE110481.

---

**Figure S1.**
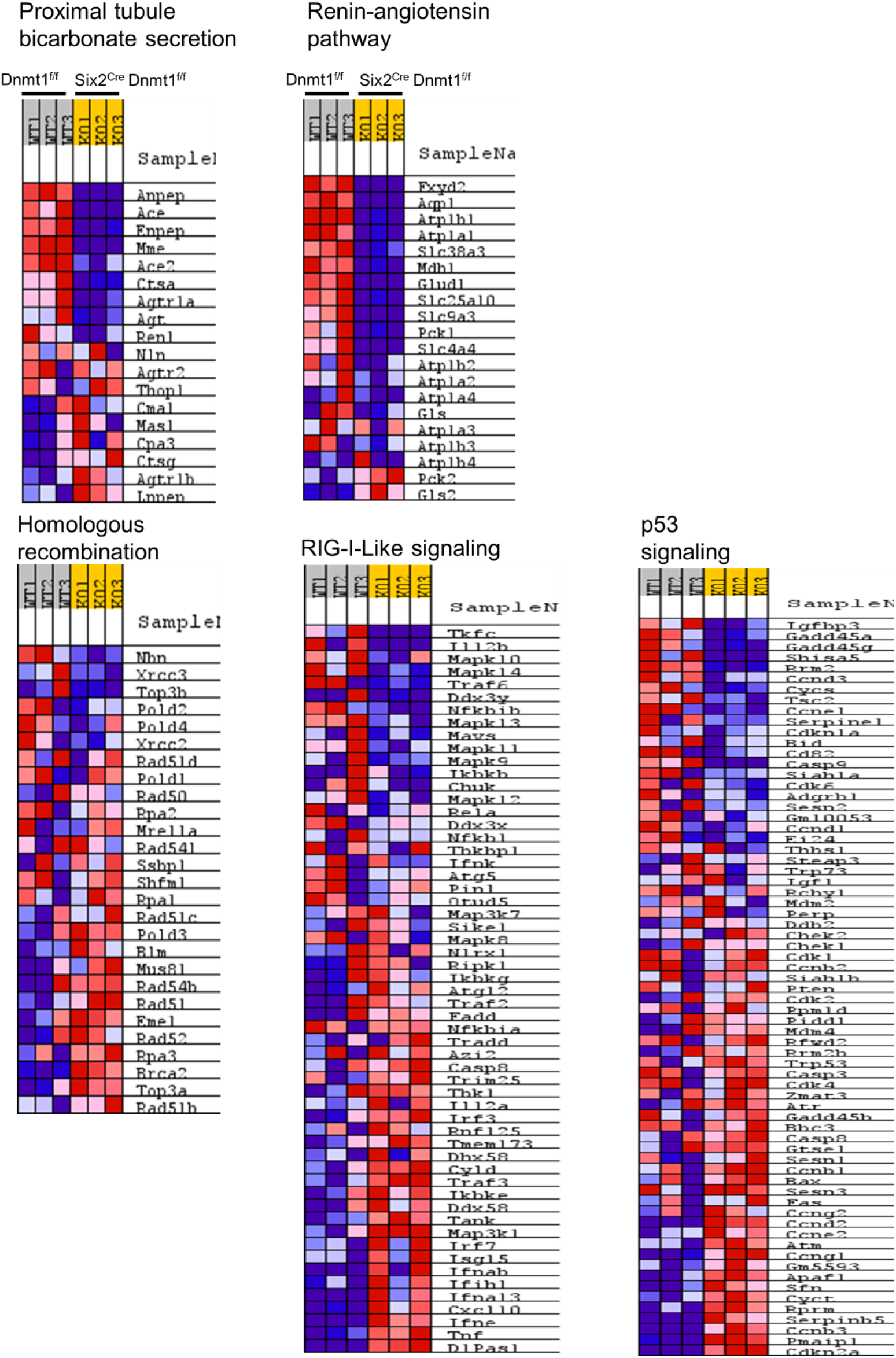
Heatmap of representative target genes showing gene expression differences. Gene set enrichment analysis of Dnmt1^f/f^ and Six2^Cre^ Dnmt1^f/f^ mice showing expression of selected pathway genes.

**Figure S2.**
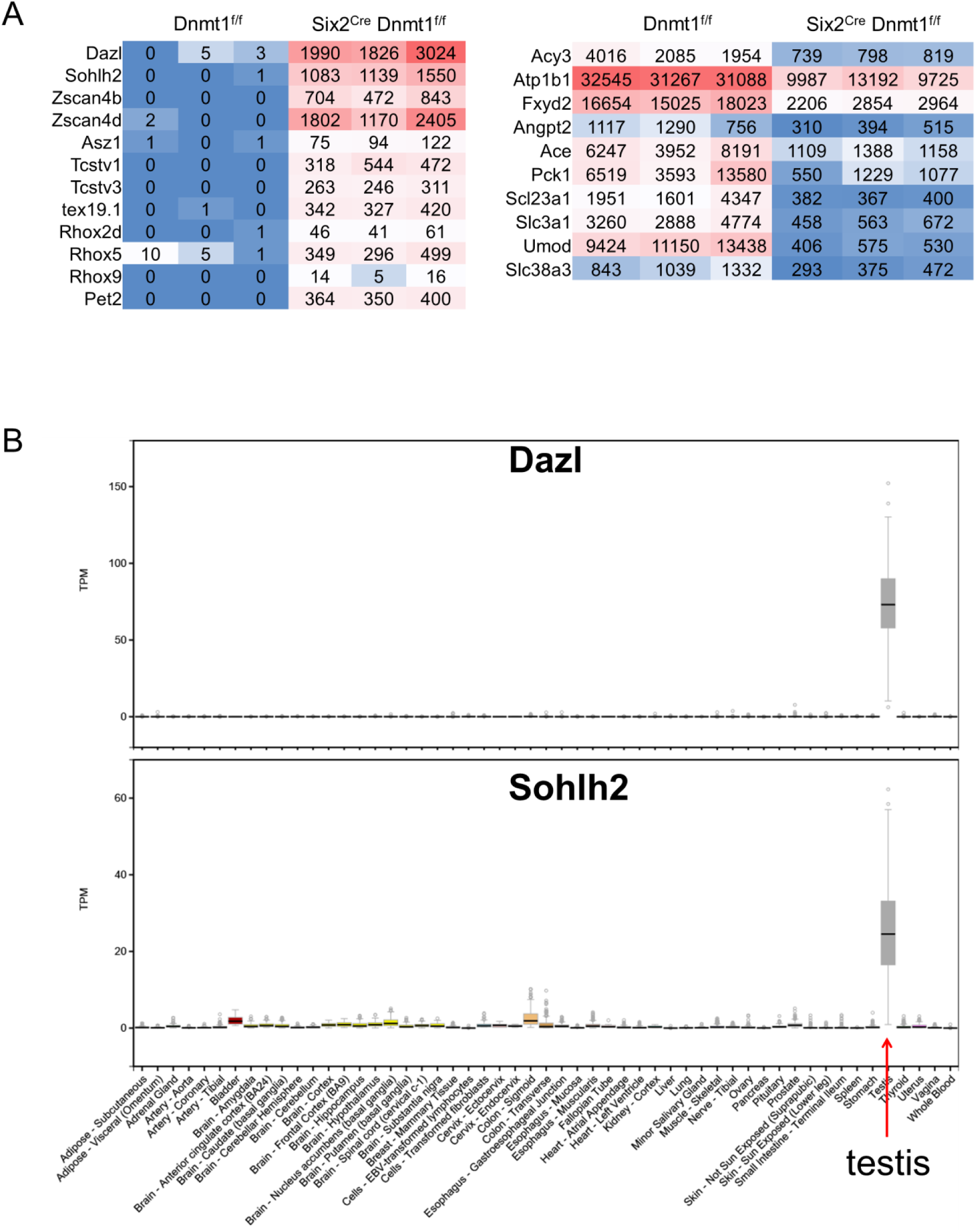
Gene expression patterns in Dnmtlt/t and Six2cre Dnmtlt/t mice. **(A)** RNAsequencing read-counts of gene expression in Dnmt1^f/f^ and Six2^Cre^ Dnmt1^f/f^ mice. **(B)** Transcript level of Dazl and Sohlh2 in different human tissue samples. Data is from GTEx Analysis Release V7.

**Figure S3.**
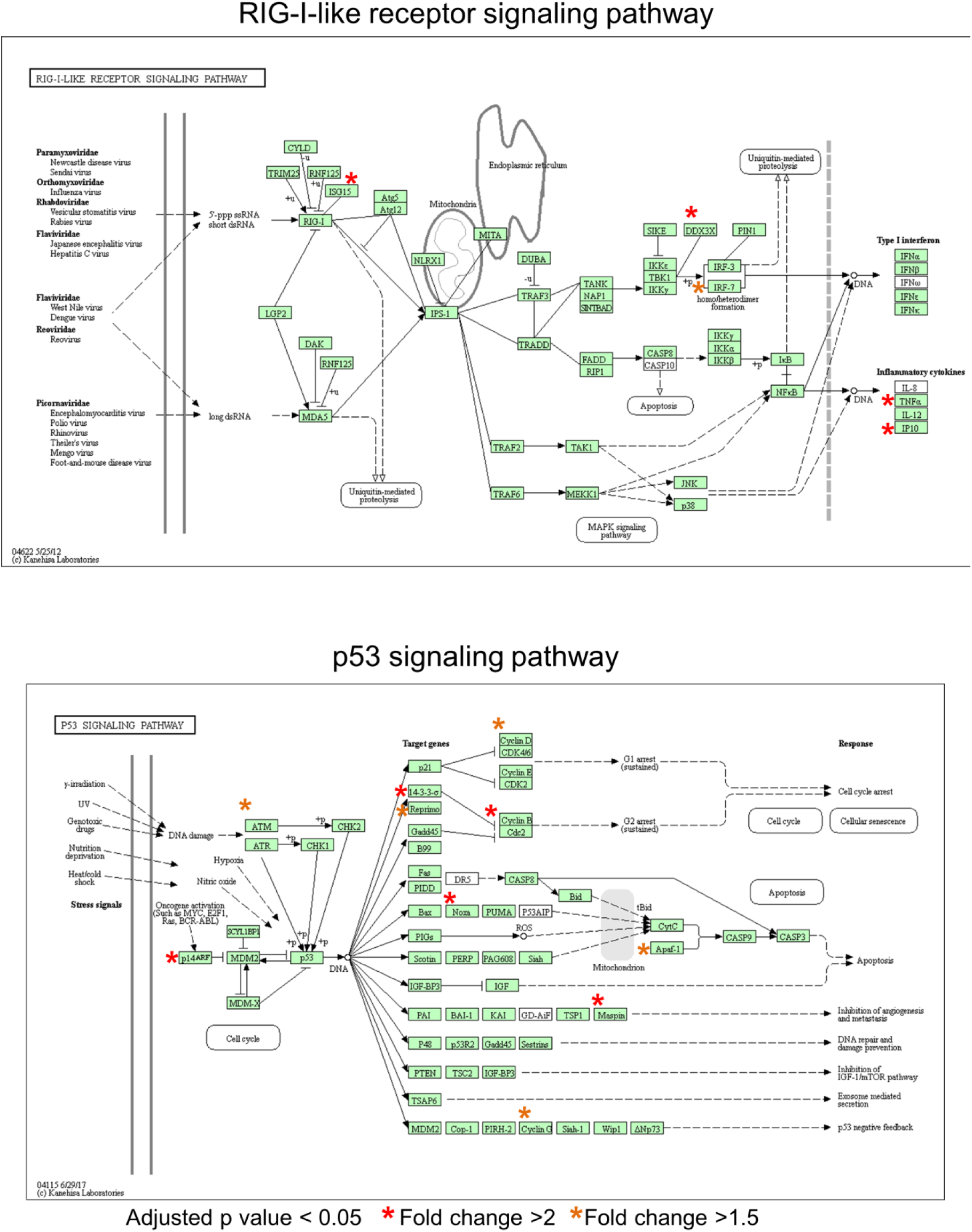
KEGG pathways for RIG-1-Iike receptor signaling pathway and P53 signaling pathway. Differentially expressed genes in between Dnmt1^f/f^ and Six2^Cre^ Dnmt1^f/f^ mice are marked by asterisks.

**Figure S4.**
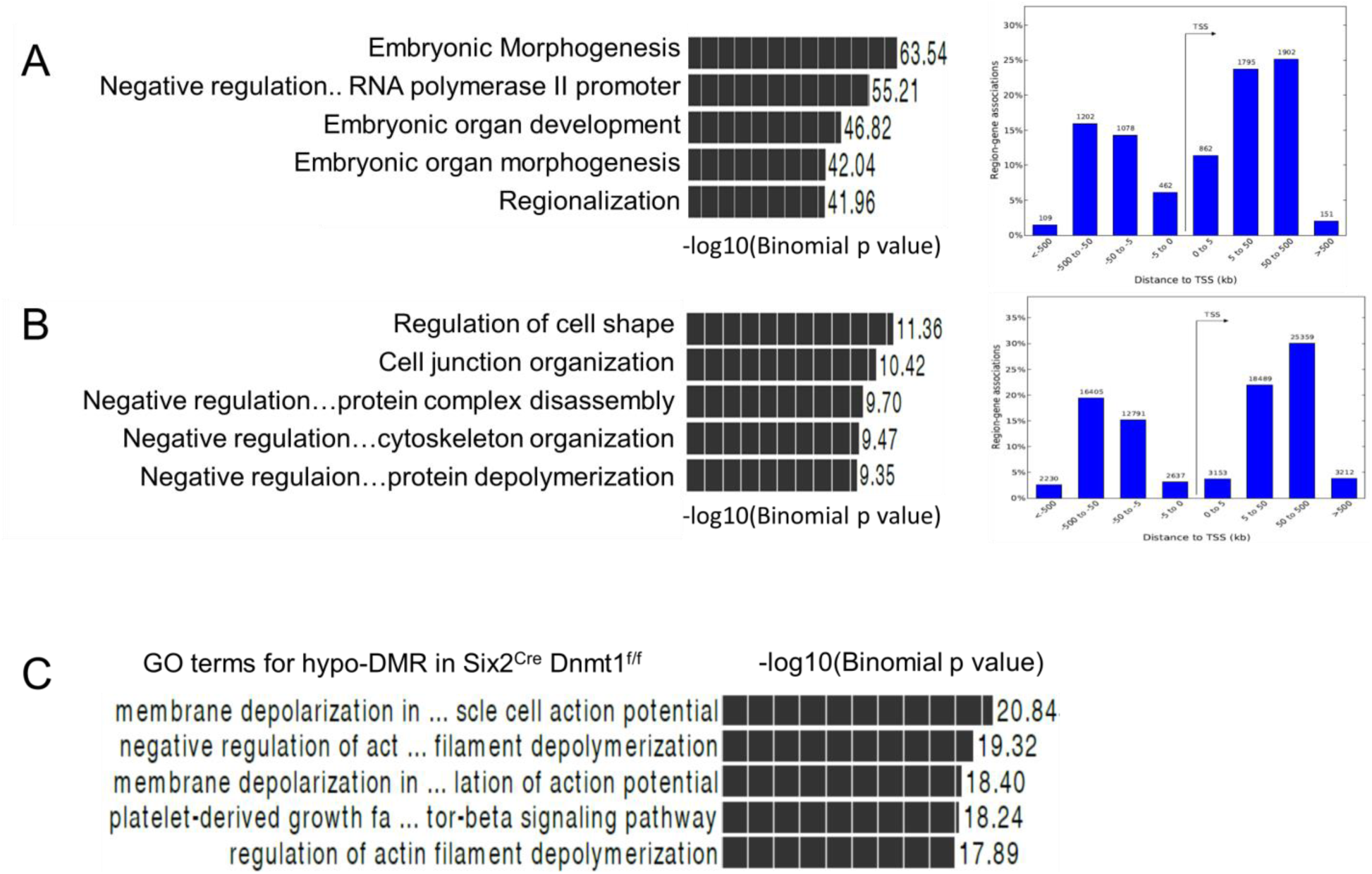
Annotation analysis of the DMRs. Left panel shows GREAT functional annotation analysis of hyper- **(A)** and hypo-DMRs **(B)** in between Dnmt1^f/f^ P0 and wild type adult mice. Right panel shows distribution of the DMRs relative to the transcription start sites. (**C**) Functional annotation (GREAT) analysis of hypo-DMRs in Six2^Cre^ Dnmt1^fl/fl^ kidneys.

**Figure S5.**
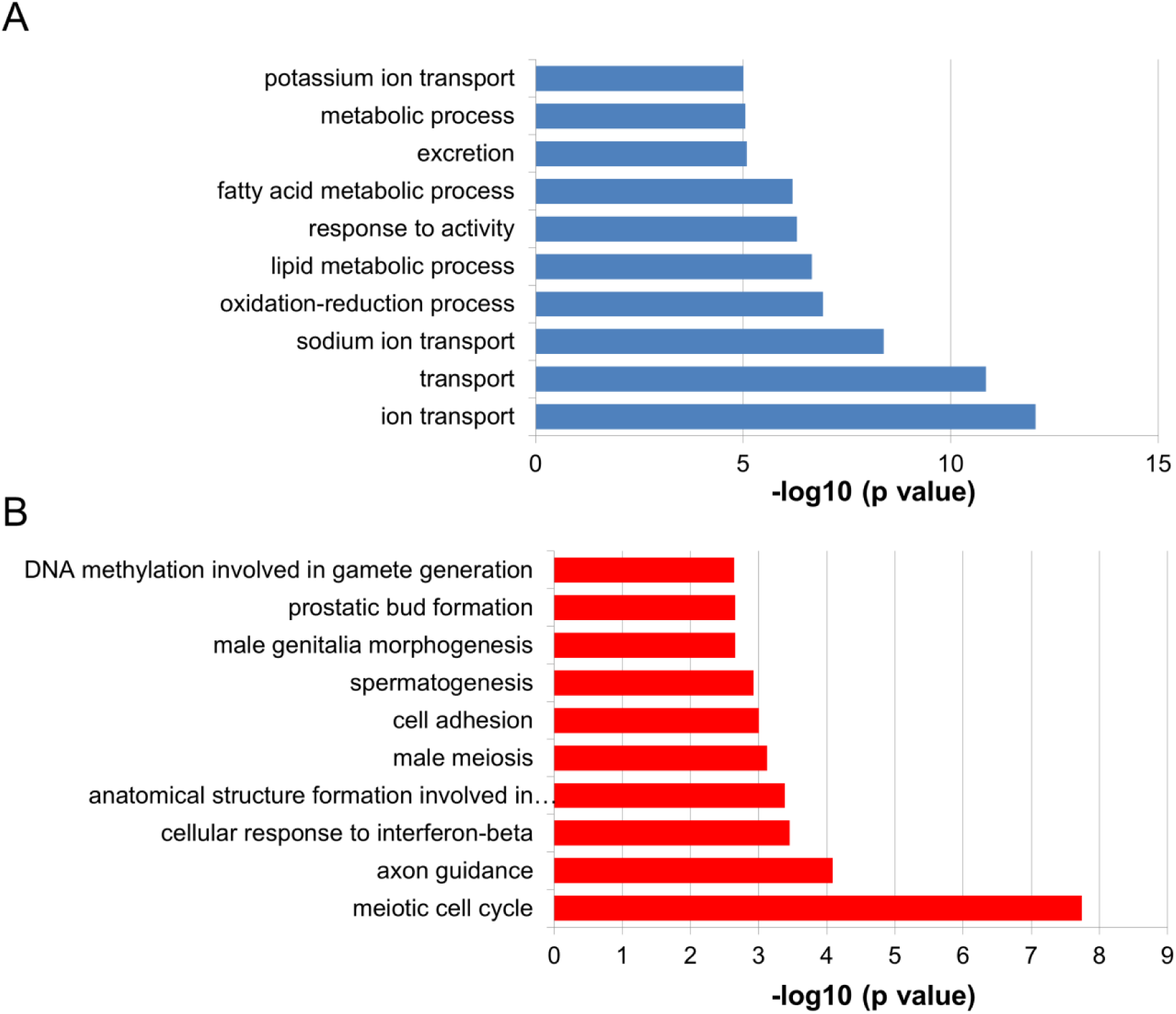
Functional annotation analysis (DAVID) Functional annotation analysis of down- **(A)** and down-regulated genes **(B)** in between Dnmt1^f/f^ and Six2^Cre^ Dnmt1^f/f^ mice.

**Figure S6.**
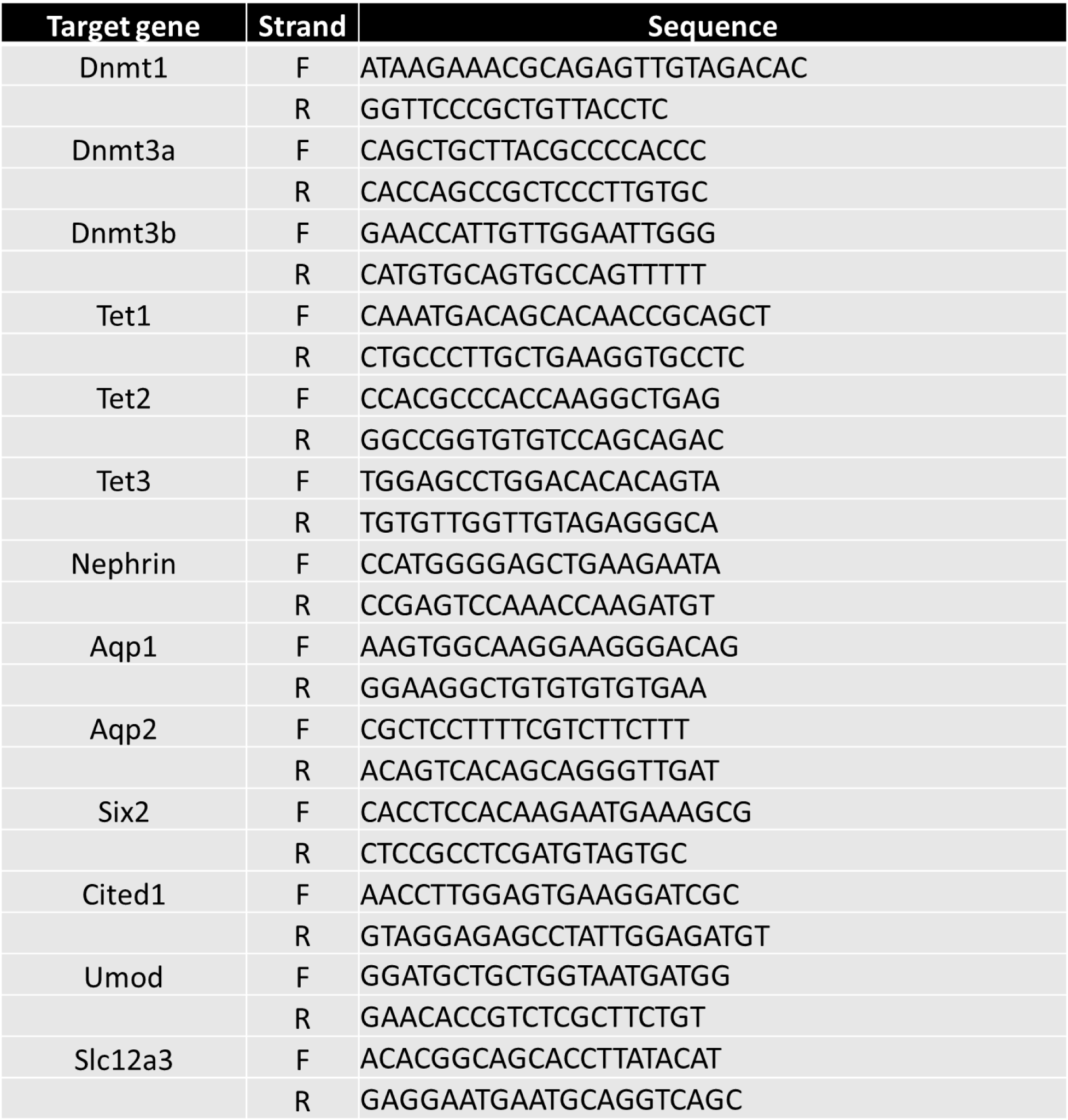
Primer sequences used in this study.

